# Hunter-gatherer admixture facilitated natural selection in Neolithic European farmers

**DOI:** 10.1101/2022.09.05.506481

**Authors:** Tom Davy, Dan Ju, Iain Mathieson, Pontus Skoglund

## Abstract

Ancient DNA has revealed multiple episodes of admixture in human prehistory during geographic expansions associated with cultural innovations. One important example is the expansion of Neolithic agricultural groups out of the Near East into Europe, and their consequent admixture with Mesolithic hunter-gatherers. Ancient genomes from this period provide an opportunity to study the role of admixture in providing new genetic variation for selection to act upon, and also to identify genomic regions that resisted hunter-gatherer introgression and may thus contribute to agricultural adaptations. We used genome-wide DNA from 728 individuals spanning Mesolithic and Neolithic Europe to infer ancestry deviations in the genomes of admixed individuals, and to test for natural selection after admixture using a new method based on testing for deviations from a genome-wide null distribution. We find that the region around the pigmentation-associated gene *SLC24A5* shows the greatest overrepresentation of Neolithic ancestry in the genome (|Z| = 3.45). In contrast, we find the greatest overrepresentation of Mesolithic local ancestry across the key immunity locus that is the Major Histocompatibility Complex (MHC; |Z| > 4) which also shows allele frequency deviations indicative of a selective sweep following admixture (p =1×10^−29^). This could reflect negative frequency dependent selection on MHC alleles common in Neolithic populations, or that Mesolithic alleles were positively selected for and facilitated adaptation by Neolithic populations to pathogens, new diets, or other environmental factors. Our results extend previous results that highlight immune function and pigmentation as targets of adaptation in more recent populations to selection processes in the Stone Age, and demonstrate that admixture facilitated selection by contributing new genetic variation.

## Introduction

Despite the evidence from studies of ancient DNA that admixture among Holocene populations is ubiquitous, less is known about how admixture provided variation for natural selection to act upon during transitional periods. Given that Holocene admixture was often associated with dramatic migrations or changes in lifestyle, we might expect an important role for adaptive introgression. Perhaps the best-studied example of ancient admixture is in the Mesolithic-Neolithic transition in Europe. As Early Neolithic groups expanded across Europe from Anatolia in the period 10,000 to 5,000 years ago^1–7^, they admixed with local Mesolithic hunter-gatherers, and by the later Middle Neolithic period derived 20-30% of their ancestry from these local groups^1,2,8,9^. The admixed Neolithic ancestry thus found itself in a new demographic, cultural, and geographic landscape. In addition to dramatic dietary changes, the Neolithic transition included an increase in population density, which together with the presence of domesticated animals has been hypothesised to lead to increased infectious disease load^10^.

Despite the potential of introgression as a source of adaptive genetic variation, examples of adaptive admixture in humans are relatively rare. Examples include American populations experiencing selection following admixture with European and African populations at immune loci^11^. Within Africa, the *DARC* ‘Duffy-null’ allele has seen at least two adaptive admixture events, spreading from the mainland to both Madagascar and the Capo Verde islands^12,13^, alongside signals at other immune genes^14,15^. There are also several examples of adaptive alleles inherited from archaic humans such as a *WARS2*-*TBX15* haplotype in Greenlandic Inuit^16^ and an allele of *EPAS1* in present-day Tibetans^17^ both introgressed from Denisovans. Studies of adaptive admixture among present-day populations are in general confined to recent admixture in the past ten or so generations, which is too short to detect anything but the strongest selection. On the other hand, signals of adaptive archaic introgression from events thousands of generations ago may have decayed with time and be difficult to detect.

Previous studies of natural selection in the European Neolithic have either compared allele frequencies or haplotype structure with other ancient and modern populations^2,18–21^. However, no study to date has attempted to assign signals of adaptive admixture to a particular ancestry close to the time of admixture. The admixture of Early Neolithic and hunter-gatherer ancestry represents an opportunity to study adaptive admixture, similar to recently admixed present-day populations, but over a much longer timescale; around a hundred generations (**Figure 1A**). A recent study identified two optimal approaches to detect adaptive admixture, based on allele frequencies and local ancestry, respectively^22^. Here, we adapt these two approaches to ancient populations with a new framework to obtain p-values from genome-wide null distributions, investigating adaptive admixture in 728 Mesolithic, Early and Middle Neolithic individuals.

**Figure 1.**
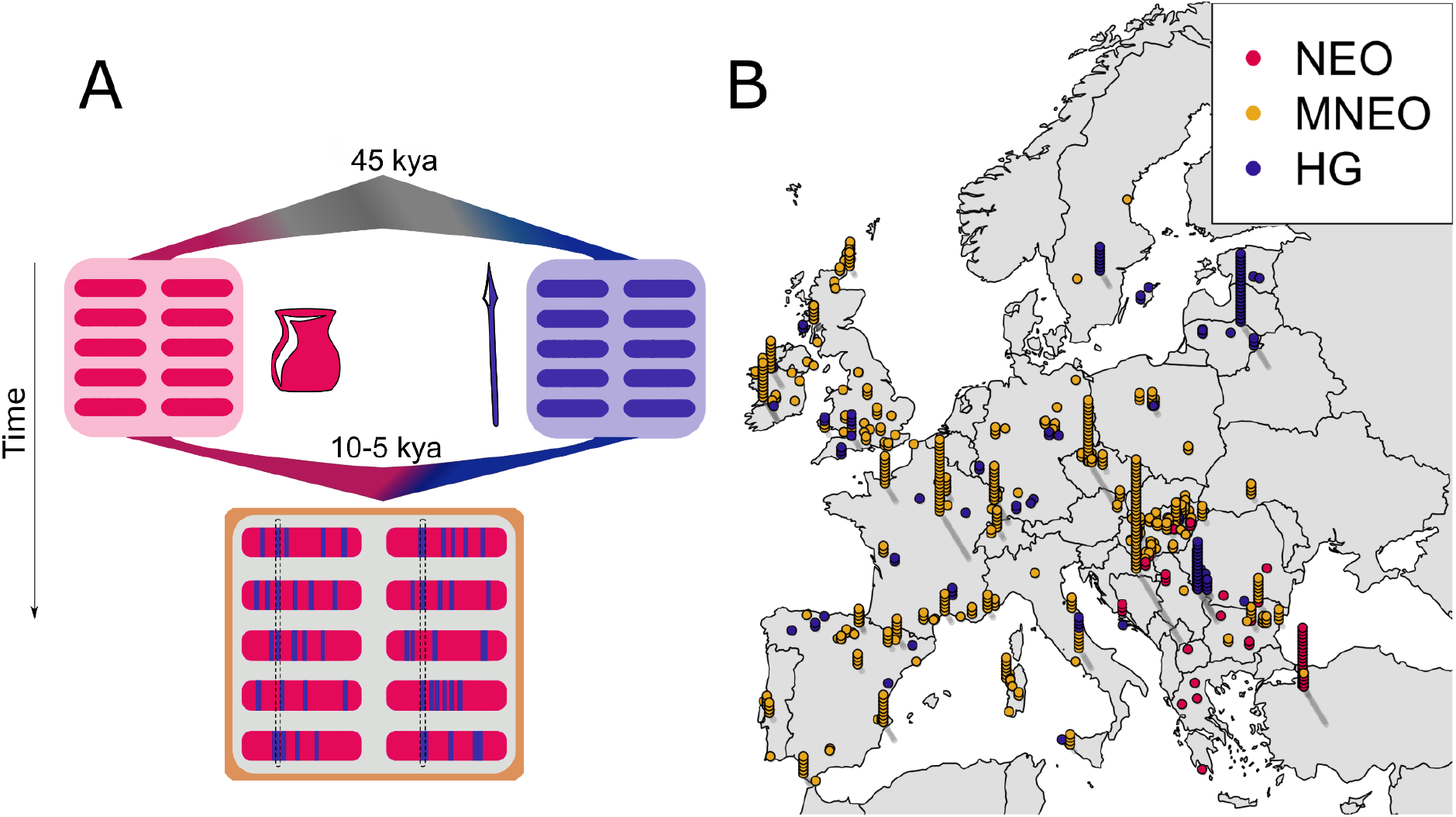
Admixture model and geographic distribution of Neolithic and Mesolithic individuals with genome-wide ancient DNA. **A)** An illustration of the genetic history of the Neolithic-Mesolithic transition in Western Eurasia. **B) ‘**Casino-plot’ of individuals included for analyses, coloured by the ancestry group for which those individuals were used in this paper. For sites with multiple samples, we stack those individuals above the reported coordinates.

## Results and Discussion

We used clustering analyses to assign individuals with genome-wide ancient DNA data from Europe and Anatolia in the past ~15,000 years to one of three groups: HG for 135 Mesolithic and Upper Palaeolithic hunter-gatherer ancestry individuals, NEO for 56 early Neolithic individuals from Anatolia and the Balkans without evidence for substantial hunter-gatherer admixture, and MNEO for 537 later Middle Neolithic individuals with substantial HG admixture. In total, our analysis contains 728 ancient individuals, spanning 7,500 years across the European continent (**Figure 1A-B; Supplementary Figure 1; Supplementary Figure 2**).

We first used an approach to find natural selection that is admixture-unaware, searching for increased differentiation as the squared allele frequency difference between populations, the *f*_2_ statistic^23^. We computed the average for each SNP and 25 SNPs flanking SNPs on each side (*i.e*. in 51 bp sliding windows), and obtain p-values by fitting a gamma distribution to a null sample of 532 approximately independent loci, separated by at least 5 million basepairs (mb). We observe no statistically significant outliers in the HG-NEO or HG-MNEO contrasts (**Supplementary Figure 3**), but in the NEO-MNEO contrast we observe a highly significant excess differentiation across the Major Histocompatibility Complex (MHC) region on chromosome 6, centred upon *HLA-DQB1* (p = 1×10^−21^) (**Supplementary Figure 3**). This suggests natural selection at the MHC in the period covered by the data from NEO and MNEO groups, although this analysis does not address whether that might be due to adaptive HG admixture.

To search for adaptive admixture, we applied a statistic based on previous work^2^ termed *F*_adm_^22^, which tests for deviations from the expected allele frequencies given the genome-wide average mixture proportions of the contributing ancestries. We again observe an excess signal across the MHC, centred upon *HLA-DQB1* (p = 1×10^−29^) (**Figure 1A**). To confirm that our findings were not driven by ascertainment bias in the 1.2M SNP panel^24^, we analysed the MHC region in 156 whole-genome shotgun sequences from 67 Mesolithic (HG), 27 Early Neolithic (NEO), and 62 Middle Neolithic Admixed (MNEO) individuals (Extended Methods), and observe a concordant peak stretching across the Class III MHC region (**Figure 1E**).

We next sought to quantify the direction of admixture by searching for deviations in local ancestry across the genome (Local Ancestry Deviation; LAD, ref. ^22^). We inferred local ancestry in 537 admixed Middle Neolithic individuals with genome-wide SNP data using *ancestryHMM^25^*, which estimates local ancestry in low-coverage genomic data using allele frequencies from two populations. We computed standard errors and Z-scores for local ancestry deviation (LAD) using an approximately independent subsample of the genome-wide distribution consisting of 555 sites separated by at least 5Mb (**Methods**).

The greatest excess of Neolithic ancestry centred on *SLC24A5* (**Figure 2C; Supplementary Figure 4; Supplementary Figure 5**), with a peak of +18.6% (|Z|=3.45). The derived *SLC24A5* allele, which is carried on the Neolithic ancestry background, is one of the two alleles which contributes most to light skin pigmentation in present-day European populations^26^. It has been previously shown to have been at relatively high frequency in the Neolithic and absent in the Mesolithic hunter-gatherers^2^, and our results show that the selection removed hunter-gatherer ancestry at this locus in later admixed Neolithic groups.

**Figure 2.**
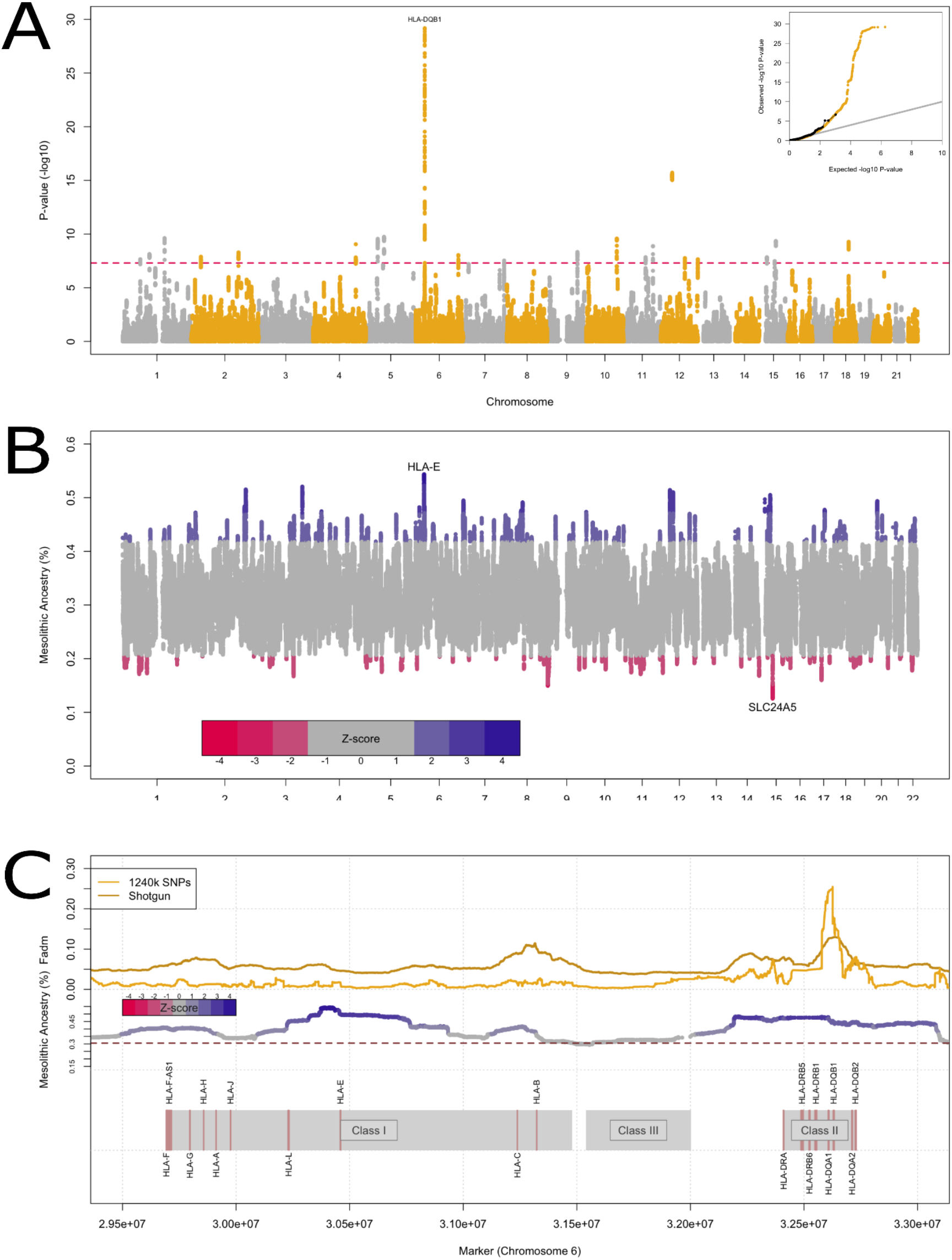
Genome-wide significant signals of adaptive admixture. **A)** Manhattan plot of p-values from the *F*_adm_ scan across the genome for deviations from expected admixed allele frequencies. Inset, quantile-quantile plot of expected and observed p-values. **B)** Local Ancestry Deviations (LAD) in the Middle-Neolithic across the genome, with top peaks of each ancestry labelled. **C)** Zoomed-in region of the MHC (chromosome 6), with statistics derived from 1240k and whole-genome shotgun data across the MHC regions I,II & III on chromosome 6.

Meanwhile, the lowest amount of Neolithic ancestry is found at the Major Histocompatibility Complex (MHC) region on chromosome 6. Within this locus, the region of highest Mesolithic ancestry is centred on *HLA-E*, with a peak excess of +23.2% (|Z|=4.34). This region of elevated Mesolithic ancestry continues as a contiguous region which extends across the Class II region of the MHC, with an average of ancestry across the entire MHC (between hg19 positions 28,477,797-33,448,354 of chromosome 6) of +9.16% (|Z|=1.82), and a secondary peak centred upon the class II region of +17.26% (|Z|=3.23) (**Figure 1E**).

In the *F*_adm_ analysis, we also find a number of genome-significant peaks outside of the MHC, though these are an order of magnitude below the MHC p-value (**Supplementary Figure 6**). *TYRP1* has been previously reported to have experienced selection in European populations^27,28^. *TLR2* is an important gene in immunity, and may also be a target of archaic adaptive admixture^29^. *MUC19* also plays a role in immunity, and was found to be under selection in Central Mexican indigenious populations^30^. *OPRM1* is documented to contribute towards skin pigmentation in Native American and East Asian populations^31,32^, while a *HRNR* haplotype linked to atopic dermatitis may have seen a selective sweep in East Asia alone^33^ *ROR1* is adjacent to *PGM1* which may have undergone selection in Mexican populations^34^. Similarly, the LAD analysis finds some genes with previous reports of selection; *HECW2* has been identified to be under selection in Khomani San^35^, a southern African group with recent hunter-gatherer lifestyle, while *PKP2* was similarly seen to be under selection in the Yoruba in west Africa in the same study.

It is also possible that adaptive admixture acted on multiple variants with small effect, spread across the genome. To test for evidence of such polygenic selection, we computed the Pearson correlation between the local ancestry deviation and the effect size for 38 traits in the UK BioBank, using genome-wide significant SNPs thinned to be approximately independent^36^ (**Figure 3**). We see significant evidence of correlation between trait scores and LAD in Skin Colour (p = 3e^−4^), consistent with the adaptive admixture around *SLC24A5*. Indeed, this signal is solely driven by two loci, with a *HERC2* variant with a skew towards the Mesolithic (Z=1.7) also contributing to a lighter level of skin pigmentation alongside *SLC24A5*. Without these two loci, there is no significant evidence of polygenic selection (P = 0.58). We also observe a weaker but significant correlation for hip size **(Figure 3, Supplementary Figure 7**).

**Figure 3.**
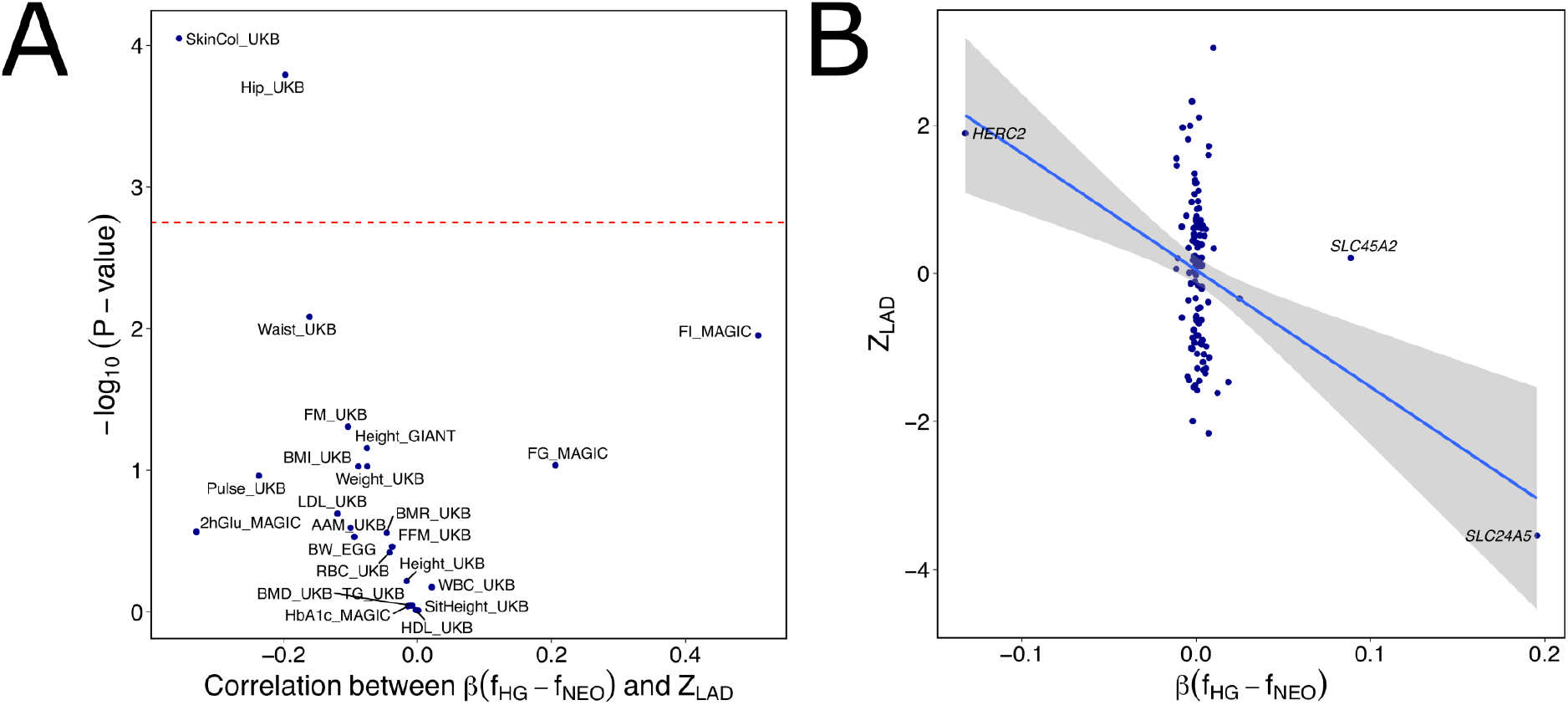
Test for polygenic adaptive admixture. **A)** Pearson correlation of polygenic traits against local ancestry. B) Correlation of LAD Z-scores with Skin Colour SNP effect size weighted by the signed allele frequency difference between the two source populations.

The Neolithic transition brought about drastic changes in demography, culture and diet, as well exposure to novel pathogens and increased potential of zoonotic disease. In admixed middle Neolithic individuals, we found excess Neolithic farmer ancestry at the pigmentation locus *SLC24A5* and excess Mesolithic hunter-gatherer ancestry at the MHC immunity locus. Previous studies also found evidence of natural selection at *SLC24A5* in European populations^26,27^ and showed that the allele was introduced into Europe in the Neolithic^2,37,38^ but our study now further demonstrates that this resulted in a removal of hunter-gatherer ancestry across the wider locus. In a similar but opposite process, the MHC locus has previously been demonstrated to have undergone selection in the ancestry of present-day Europe^239^ and specifically in Neolithic Europe^18^. Here, we obtain further robust results for selection at the MHC locus corrected for multiple testing, and demonstrate that this process specifically increased hunter-gatherer ancestry at the locus.

In contrast to *SLC24A5*, the second high-effect pigmentation variant, in *HERC*, displays an excess of Mesolithic ancestry (+17.23%, |Z| = ~3.11). Together with the third high-effect pigmentation variant at *SLC45A2*, which arrived in Europe via later expansions from the steppe, selection on pigmentation in Europe thus targeted variants from each of the three major ancestral populations^9^. This highlights the prominent role of admixture in the evolution of skin pigmentation in Western Eurasia. That this signal is not found in the allele-frequency based analysis with *F*_adm_ can likely be attributed to the small absolute change in allele frequency between our Neolithic populations, confirming recent demonstrations that local ancestry can in some cases be more powerful than allele frequency analysis for detecting selection in admixed popuations^22^.

Evidence of selection on Mesolithic ancestry across the MHC locus, highlights its role in facilitating adaptation in immunity during the Neolithic transition in Europe. One hypothesis is that this reflects the fact that Neolithic populations were expanding into environments containing pathogens to which Mesolithic populations had already adapted. This is contrary to the idea that the pathogen load in Neolithic populations was solely driven by increased population density and proximity to zoonotic vectors via animal husbandry. While examples of putative adaptive admixture involving the MHC have been previously described^11,15,40^, a clear link between the alleles under selection within this region and a specific pathogen has not been previously observed.

Another possibility is that this adaptation reflects negative frequency-dependent selection^41^, for example because pathogens adapt to the host population’s immunity genes, with a bias towards more common variants. Rare alleles would thus confer a fitness advantage, until they become common enough that other, now-rarer, alleles have higher fitness. Under this model, HLA alleles unseen by a given pathogen will attain a higher fitness initially following admixture owing to their novelty for the pathogen^42^. Thus, the selection on the minor Mesolithic ancestry could simply reflect admixture introducing rarer variants into Neolithic populations. Future studies, including whole-genome shotgun data in tandem with improved functional annotation, may shed further light on this adaptive process.

## Acknowledgements

We thank Leo Speidel for helpful comments. I.M was funded by the NIGMS ([R35GM133708] to I.M.). P.S. was supported by the EMBO Young Investigator Programme, the Vallee Foundation, the European Research Council (grant no. 852558), the Wellcome Trust (217223/Z/19/Z), and Francis Crick Institute core funding (FC001595) from Cancer Research UK, the UK Medical Research Council, and the Wellcome Trust. This research was funded in whole, or in part, by the Wellcome Trust (FC001595 and 217223/Z/19/Z). For the purpose of Open Access, the author has applied a CC BY public copyright licence to any Author Accepted Manuscript version arising from this submission. The content is solely the responsibility of the authors and does not necessarily represent the official views of the funding agencies.

## Experimental Procedures

### Data preparation

We first used clustering approaches on a large set of previously published^1–9,37,38,43–66^ individuals to identify those individuals exclusively or almost exclusively HG (Mesolithic, Western or Siberian Hunter-Gatherers [WHG, SHG]) or NEO ancestry (Early Neolithic, which we identify using Neolithic Anatolia as a baseline). We downloaded v50 of 10,391 individuals (3,589 ancient) from the Reich lab https://reich.hms.harvard.edu/downloadable-genotypes-present-day-and-ancient-dna-data-compiled-published-papers, accessed July 19^th^ 2022 for the 1240K set of SNPs. We ran a first set of clustering with ADMIXTURE from K=3 to K=11, conditioned on European individuals dated to >5kya excluding siblings, parents, and duplicates. We then further filtered on age, setting a max age for HG, NEO and MNEO of 12kya, 8.5kya and 8kya respectively, retaining 728 individuals. Using PLINK, we then make a subset of files containing only individuals fitting into either Mesolithic (EHG+WHG) or admixed Neolithic ancestry, alongside a geographically and temporally conservative Neolithic cluster.

### Detecting selective sweeps

We filtered the full set of SNPs by conditioning on observing at least 10 non-missing pseudohaploid genotypes in earch population. We utilised the F_adm_ statistic, i.e. (F_MNEO_ – E(F_MNEO_))^2^ / (1 – E(F_MNEO_)^2^), where E(F_MNEO_), where F is the frequency of a given SNP and E(F_MNEO_) is an expectation of allele frequency derived by the contributing ancestry proportions weighted by allele frequency (F_MESO_Anc_MESO_ + F_NEO_Anc_NEO_). In all cases, the given statistic is calculated on a per-SNP basis before sliding-window is applied across the genome with a step size of 1. We derive the *f*_2_ analyses in an identical manner. We annotate genes using the gencode v39liftover37 annotation files.

For each analysis, we then draw a null distribution from this sliding-window whole-genome distribution, ensuring that each sliding-window datum that contributes to the null distribution is sampled with SNPs that are, as a group, no less than 5Mb away from the previous sample. A gamma distribution is then fitted to this null distribution using the R package *fitdistrplus v1.1-3* ^67^ with the flags ““gamma”, method = “mle”, keepdata = T”. We then derive p-values for the genome-wide distribution from this null-fitted gamma distribution..

### Confirmation of HLA signal in whole-genome shotgun data

To confirm the presence of the HLA signal in the whole-genome shotgun data, we collected a set of 81 individuals used in the original analyses which had whole-genome shotgun genomes, and added 74 unpublished genomes available pre-publication under the Ft. Lauderdale principles (https://reich.hms.harvard.edu/ancient-genome-diversity-project). We thus only use this data for local analyses of the MHC region. Exclusion of the pre-publication data results in too few NEO source genomes for reliable inference, and so we do not report genome-wide p-values. We again partition out our target populations using appropriate populations within this dataset. We filter the region initially on sites with more than 3 allele counts seen within the whole dataset, and a minimum of 30 sites seen in the Neolithic population, resulting in 11,086,998 sites after filtering. We apply a sliding window of 593 snps on the mixed-weight statistic in a similar way to the 1240k analysis, using the same weighting for ancestry contribution to SNP frequency expectation. We choose this window size to approximate the same mean genetic distance per window as in the 1240k analysis with 51 snp windows.

### Detection of Local Ancestry Outliers

To analyse biases in local ancestry, we progress the same panel of 728 individuals (Supplementary Table 1) to analysis with *AncestryHMM* ^25^. We derive frequency counts for both the contributing ancestries (HG, NEO) and similarly obtain this for each individual MNEO sample via plink. We then run Ancestry HMM with a prior of 30% HG and 70% Neolithic ancestry, and an *N*_e_ of 10,000.

### Detection for biased inheritance of polygenic trait alleles

As in ref. ^36^, we start with 28 quantitative traits of interest, subsetting SNPs P < 1e^−8^ overlapping the 1240k SNP array. We then iteratively prune through subsetting the smallest p-value and removing all other associations within 250kb. Departing from Mathieson & Terhost, we weight these pruned values by the difference in allele frequency between HG and NEO and sum these across the genome for each trait. and look for correlation to the LAD derived in the local ancestry analysis via a Pearson correlation test.

## Supplementary Figures

**Supplementary Figure 1.**
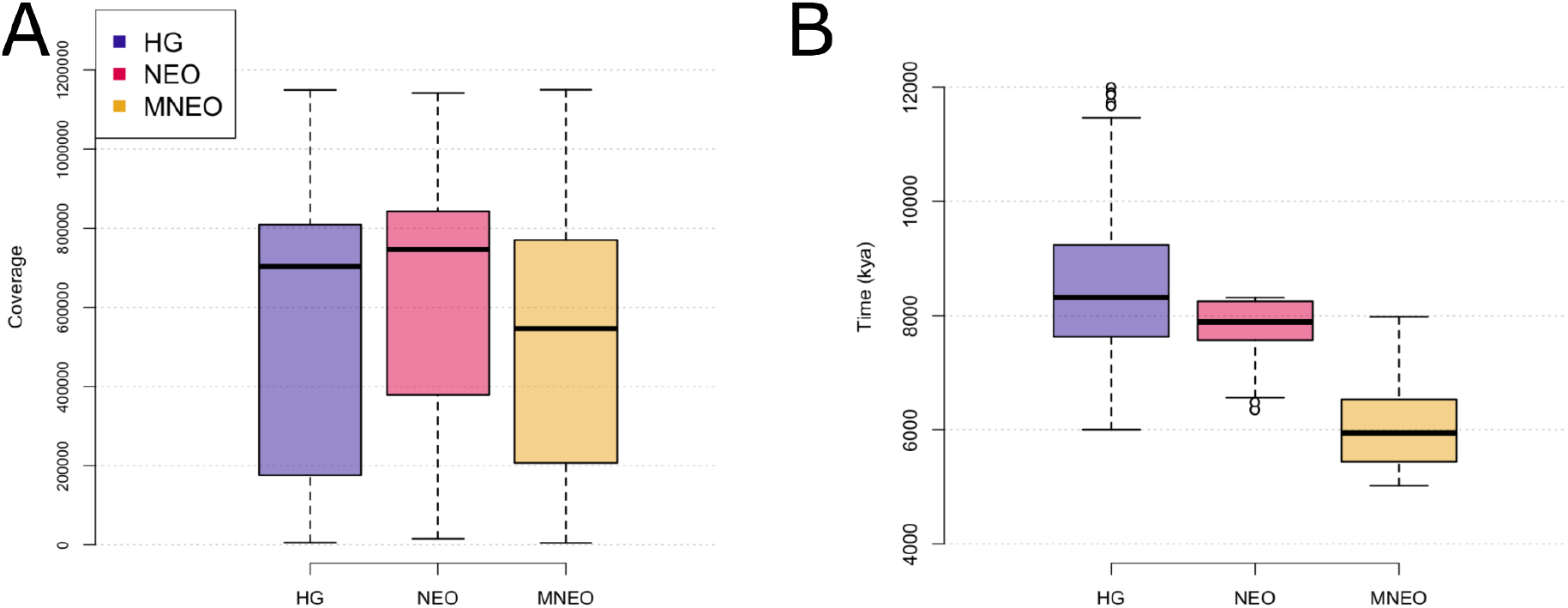
Metadata of individuals used in selection scan. A) Coverage in each population, as the number of 1240k sites covered in each individual per population. **B)** Age of individuals in the 1240k selection scan.

**Supplementary Figure 2.**
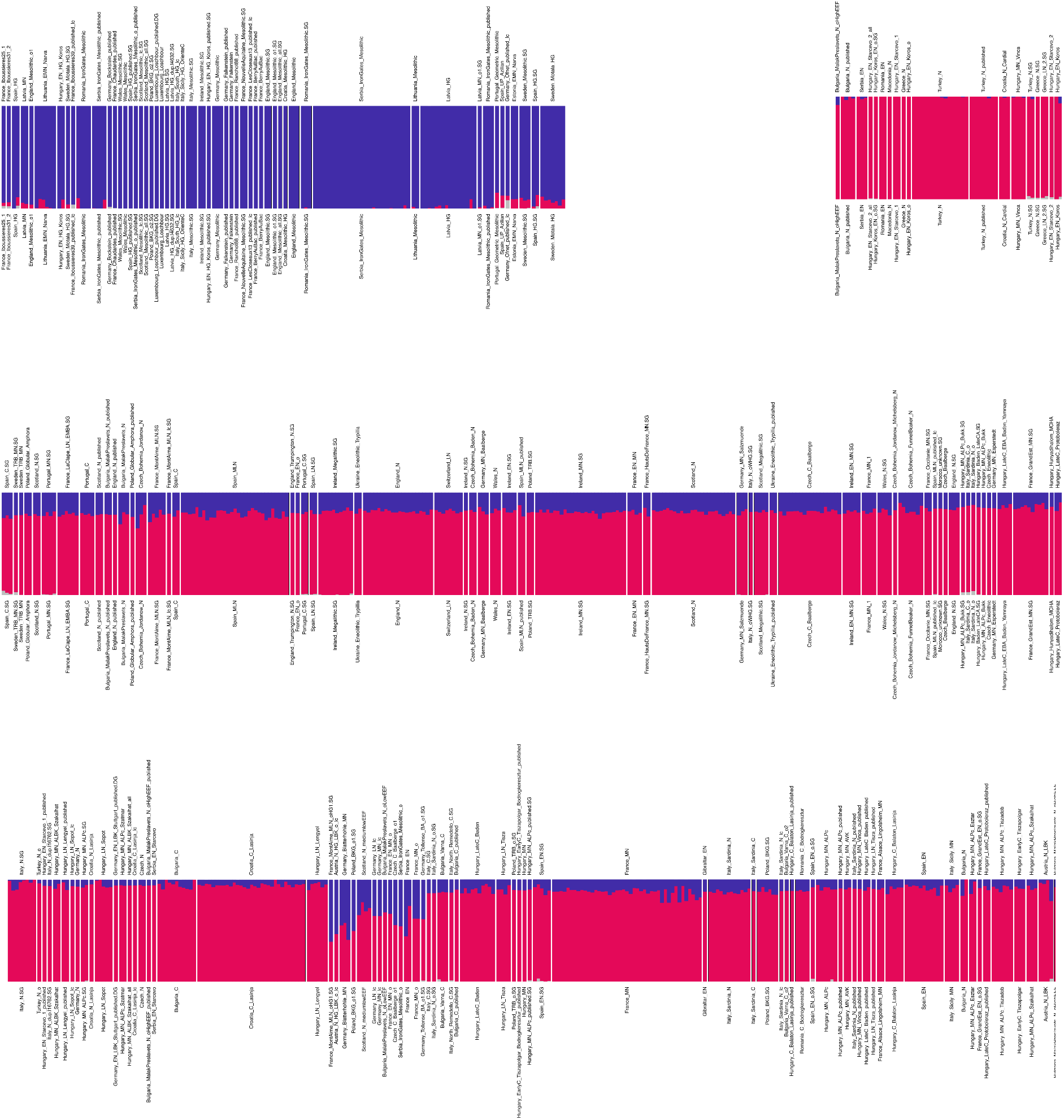
Clustering results of 728 HG, NEO, and MNEO individuals alongside 40 individuals from the MSL 1000 genomes panel (the latter not shown) obtained using ADMIXTURE with K=3 (Methods).

**Supplementary Figure 3.**
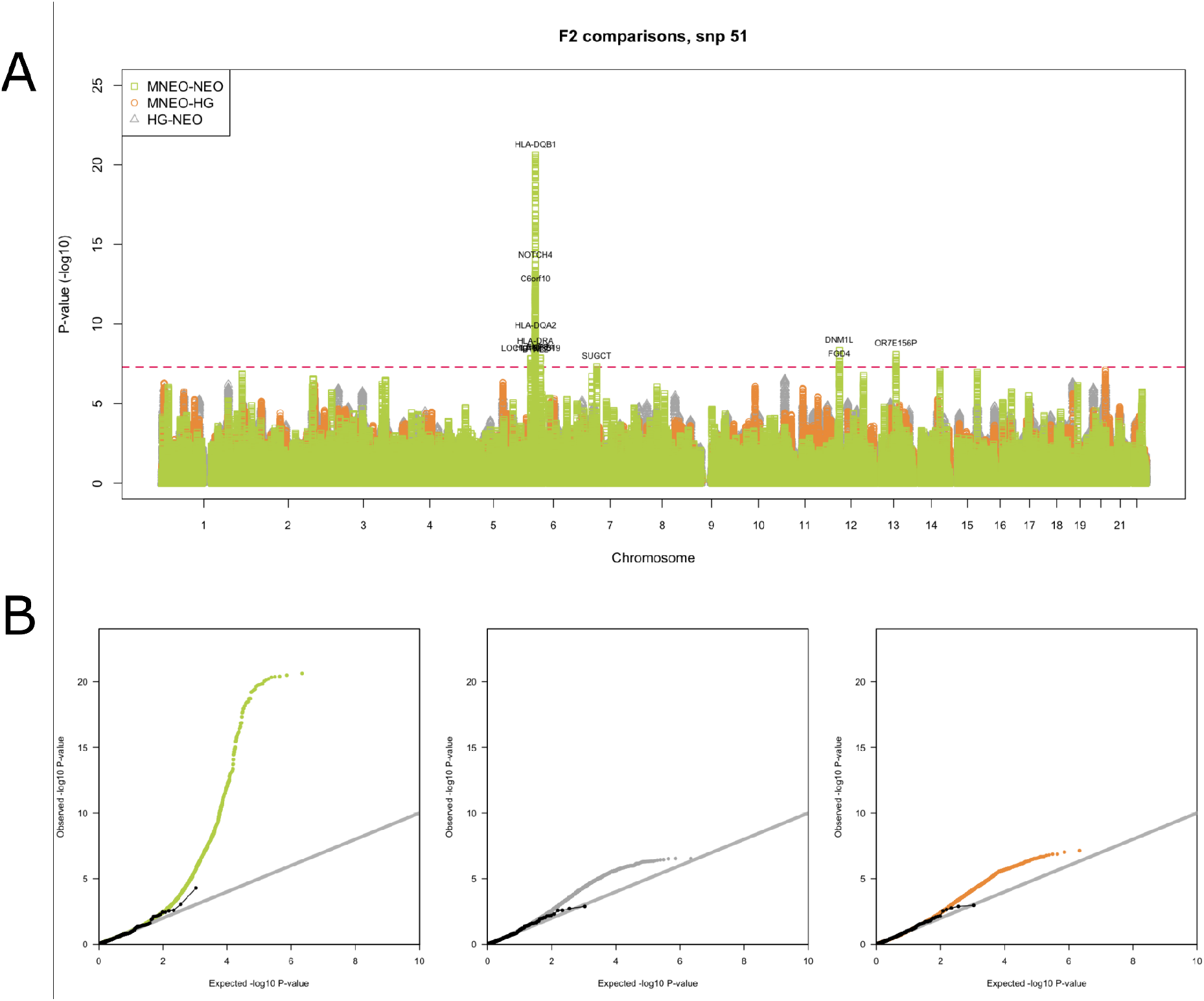
*f*_2_ pairwise comparisons of ancient populations. A) Manhattan plot of p-values for each pairwise comparison. B) Quantile-quantile plots for each pairwise comparison.

**Supplementary Figure 4.**
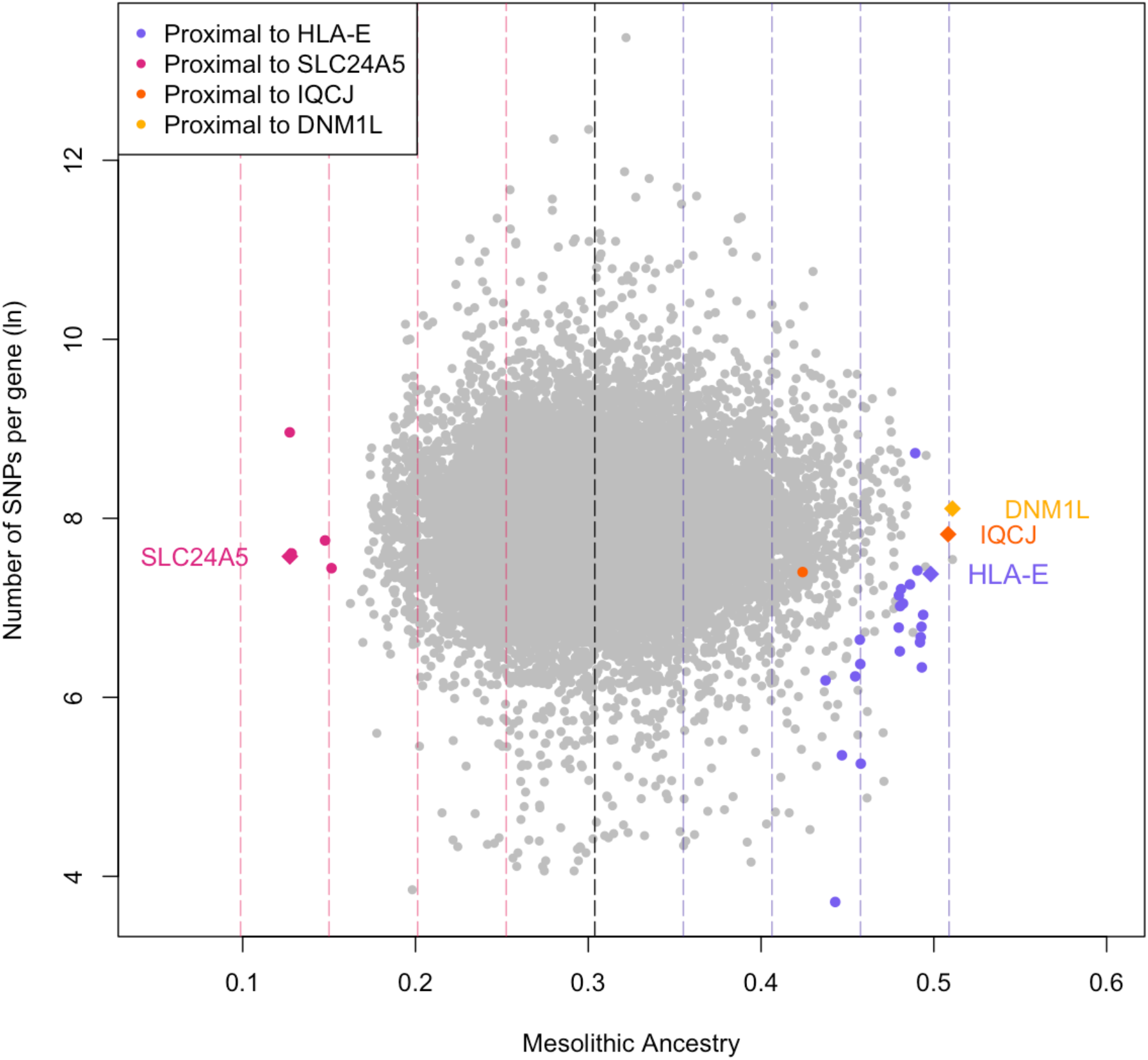
Mesolithic ancestry per gene. Proximal genes are defined as being within 300kb from the start or end of the gene. Dotted lines represent standard deviations from the mean.

**Supplementary Figure 5.**
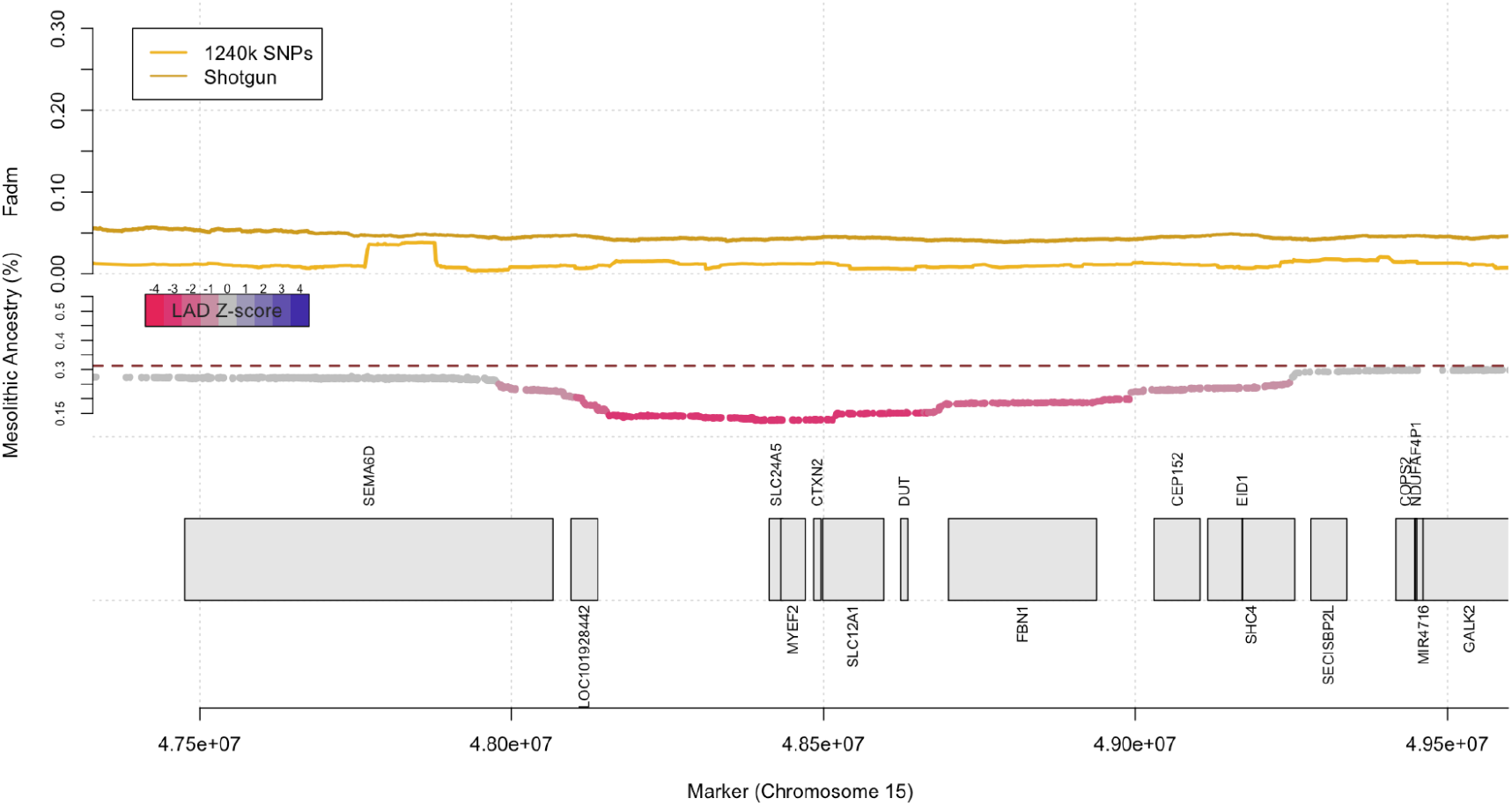
Zoomed-in region of chromosome 11, with adaptive admixture statistics derived from 1240k and Shotgun data alongside local ancestry signal throughout the MHC regions I,II & III on chromosome 15.

**Supplementary Figure 6.**
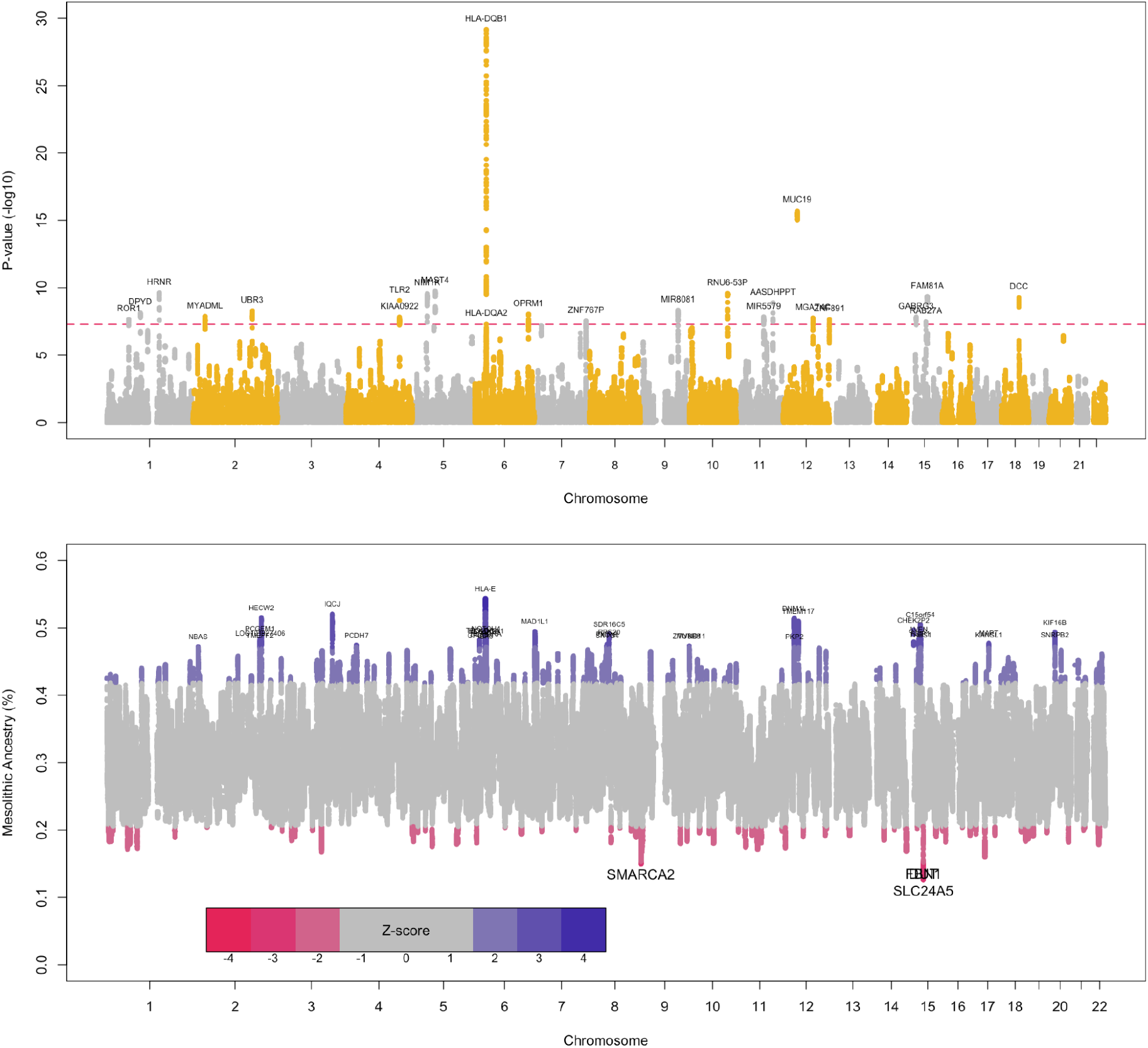
Fully annotated versions of adaptive admixture test Manhattan plots as seen in figure 2. A) F_adm_ **B**) Local Ancestry Deviation.

**Supplementary Figure 7.**
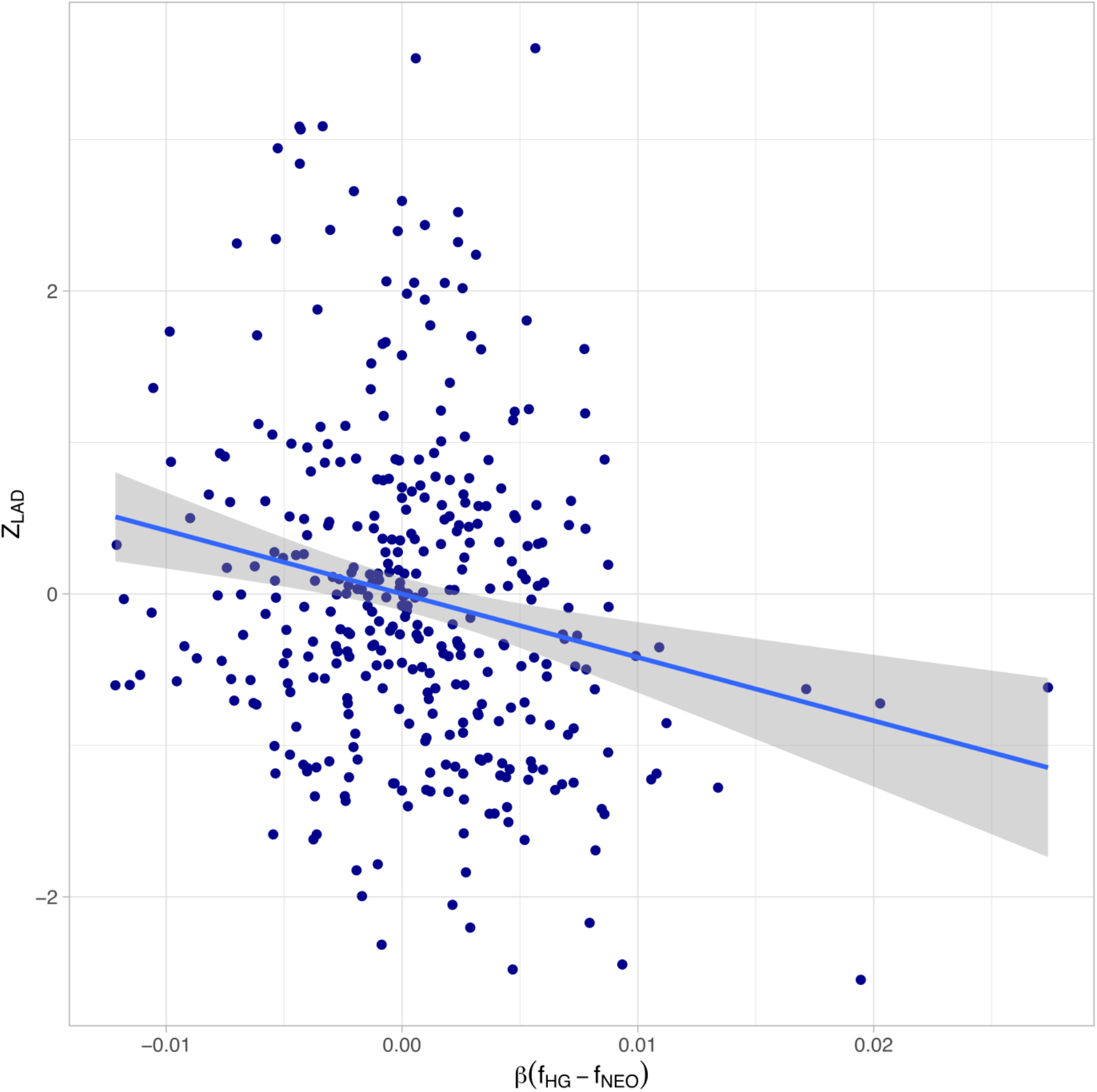
Correlation of LAD Z-scores with hip circumference SNP effect size weighted by the signed allele frequency difference between the two source populations.

## References

1. Haak, W., Lazaridis, I., Patterson, N., Rohland, N., Mallick, S., Llamas, B., Brandt, G., Nordenfelt, S., Harney, E., Stewardson, K., et al. (2015). Massive migration from the steppe was a source for Indo-European languages in Europe. Nature 522, 207.

2. Mathieson, I., Lazaridis, I., Rohland, N., Mallick, S., Patterson, N., Roodenberg, S.A., Harney, E., Stewardson, K., Fernandes, D., Novak, M., et al. (2015). Genome-wide patterns of selection in 230 ancient Eurasians. Nature 528, 499–503.

3. Hofmanová, Z., Kreutzer, S., Hellenthal, G., Sell, C., Diekmann, Y., Díez-del-Molino, D., van Dorp, L., López, S., Kousathanas, A., Link, V., et al. (2016). Early farmers from across Europe directly descended from Neolithic Aegeans. Proceedings of the National Academy of Sciences 113, 6886–6891.

4. Cassidy, L.M., Martiniano, R., Murphy, E.M., Teasdale, M.D., Mallory, J., Hartwell, B., and Bradley, D.G. (2016). Neolithic and Bronze Age migration to Ireland and establishment of the insular Atlantic genome. Proceedings of the National Academy of Sciences 113, 368–373.

5. Kılınç, G.M., Koptekin, D., Atakuman, Ç., Sümer, A.P., Dönertaş, H.M., Yaka, R., Bilgin, C.C., Büyükkarakaya, A.M., Baird, D., Altınışık, E., et al. (2017). Archaeogenomic analysis of the first steps of Neolithization in Anatolia and the Aegean. Proc. Biol. Sci. 284.

6. Omrak, A., Günther, T., Valdiosera, C., Svensson, E.M., Malmström, H., Kiesewetter, H., Aylward, W., Storå, J., Jakobsson, M., and Götherström, A. (2015). Genomic Evidence Establishes Anatolia as the Source of the European Neolithic Gene Pool. Curr. Biol. 26, 270–275.

7. Kılınç, G.M., Omrak, A., Özer, F., Günther, T., Büyükkarakaya, A.M., Bıçakçı, E., Baird, D., Dönertaş, H.M., Ghalichi, A., Yaka, R., et al. (2016). The Demographic Development of the First Farmers in Anatolia. Curr. Biol. 26, 2659–2666.

8. Lipson, M., Szécsényi-Nagy, A., Mallick, S., Pósa, A., Stégmár, B., Keerl, V., Rohland, N., Stewardson, K., Ferry, M., Michel, M., et al. (2017). Parallel palaeogenomic transects reveal complex genetic history of early European farmers. Nature 551, 368–372.

9. Lazaridis, I., Patterson, N., Mittnik, A., Renaud, G., Mallick, S., Kirsanow, K., Sudmant, P.H., Schraiber, J.G., Castellano, S., Lipson, M., et al. (2014). Ancient human genomes suggest three ancestral populations for present-day Europeans. Nature 513, 409–413.

10. Barrett, R., Kuzawa, C.W., McDade, T., and Armelagos, G.J. (1998). EMERGING AND RE-EMERGING INFECTIOUS DISEASES: The Third Epidemiologic Transition. Annual Review of Anthropology 27, 247–271.

11. Mendoza-Revilla, J., Chacón-Duque, J.C., Fuentes-Guajardo, M., Ormond, L., Wang, K., Hurtado, M., Villegas, V., Granja, V., Acuña-Alonzo, V., Jaramillo, C., et al. (2021). Disentangling signatures of selection before and after European colonization in Latin Americans. bioRxiv, 2021.11.15.467418.

12. Hamid, I., Korunes, K.L., Beleza, S., and Goldberg, A. (2021). Rapid adaptation to malaria facilitated by admixture in the human population of Cabo Verde. Elife 10.

13. Hodgson, J.A., Pickrell, J.K., Pearson, L.N., Quillen, E.E., Prista, A., Rocha, J., Soodyall, H., Shriver, M.D., and Perry, G.H. (2014). Natural selection for the Duffy-null allele in the recently admixed people of Madagascar. Proc. Biol. Sci. 281, 20140930.

14. Lopez, M., Choin, J., Sikora, M., Siddle, K., Harmant, C., Costa, H.A., Silvert, M., Mouguiama-Daouda, P., Hombert, J.-M., Froment, A., et al. (2019). Genomic Evidence for Local Adaptation of Hunter-Gatherers to the African Rainforest. Curr. Biol. 29, 2926–2935.e4.

15. Patin, E., Lopez, M., Grollemund, R., Verdu, P., Harmant, C., Quach, H., Laval, G., Perry, G.H., Barreiro, L.B., Froment, A., et al. (2017). Dispersals and genetic adaptation of Bantu-speaking populations in Africa and North America. Science 356, 543–546.

16. Racimo, F., Gokhman, D., Fumagalli, M., Ko, A., Hansen, T., Moltke, I., Albrechtsen, A., Carmel, L., Huerta-Sánchez, E., and Nielsen, R. (2017). Archaic Adaptive Introgression in TBX15/WARS2. Mol. Biol. Evol. 34, 509–524.

17. Huerta-Sánchez, E., Jin, X., Asan, Bianba, Z., Peter, B.M., Vinckenbosch, N., Liang, Y., Yi, X., He, M., Somel, M., et al. (2014). Altitude adaptation in Tibetans caused by introgression of Denisovan-like DNA. Nature 512, 194–197.

18. Childebayeva, A., Rohrlach, A.B., Barquera, R., Rivollat, M., Aron, F., Szolek, A., Kohlbacher, O., Nicklisch, N., Alt, K.W., Gronenborn, D., et al. (2022). Population Genetics and Signatures of Selection in Early Neolithic European Farmers. Molecular Biology and Evolution 39.

19. Ju, D., and Mathieson, I. (2021). The evolution of skin pigmentation-associated variation in West Eurasia. Proc. Natl. Acad. Sci. U. S. A. 118.

20. Allentoft, M.E., Sikora, M., Refoyo-Martínez, A., Irving-Pease, E.K., Fischer, A., Barrie, W., Ingason, A., Stenderup, J., Sjögren, K.-G., Pearson, A., et al. (2022). Population Genomics of Stone Age Eurasia. bioRxiv, 2022.05.04.490594.

21. Kerner, G., Neehus, A.-L., Abel, L., Casanova, J.-L., Patin, E., Laval, G., and Quintana-Murci, L. (2022). Genetic adaptation to pathogens and increased risk of inflammatory disorders in post-Neolithic Europe. bioRxiv, 2022.07.02.498543.

22. Cuadros-Espinoza, S., Laval, G., Quintana-Murci, L., and Patin, E. (2022). The genomic signatures of natural selection in admixed human populations. Am. J. Hum. Genet.

23. Reich, D., Thangaraj, K., Patterson, N., Price, A.L., and Singh, L. (2009). Reconstructing Indian population history. Nature 461, 489–494.

24. Lachance, J., and Tishkoff, S.A. (2013). SNP ascertainment bias in population genetic analyses: why it is important, and how to correct it. Bioessays 35, 780–786.

25. Corbett-Detig, R., and Nielsen, R. (2017). A Hidden Markov Model Approach for Simultaneously Estimating Local Ancestry and Admixture Time Using Next Generation Sequence Data in Samples of Arbitrary Ploidy. PLoS Genet. 13, e1006529.

26. Lamason, R.L., Mohideen, M.-A.P.K., Mest, J.R., Wong, A.C., Norton, H.L., Aros, M.C., Jurynec, M.J., Mao, X., Humphreville, V.R., Humbert, J.E., et al. (2005). SLC24A5, a putative cation exchanger, affects pigmentation in zebrafish and humans. Science 310, 1782–1786.

27. Voight, B.F., Kudaravalli, S., Wen, X., and Pritchard, J.K. (2006). A map of recent positive selection in the human genome. PLoS Biol. 4, e72.

28. Pickrell, J.K., Coop, G., Novembre, J., Kudaravalli, S., Li, J.Z., Absher, D., Srinivasan, B.S., Barsh, G.S., Myers, R.M., Feldman, M.W., et al. (2009). Signals of recent positive selection in a worldwide sample of human populations. Genome Res. 19, 826–837.

29. Enard, D., and Petrov, D.A. (2018). Evidence that RNA Viruses Drove Adaptive Introgression between Neanderthals and Modern Humans. Cell 175, 360–371.e13.

30. Reynolds, A.W., Mata-Míguez, J., Miró-Herrans, A., Briggs-Cloud, M., Sylestine, A., Barajas-Olmos, F., Garcia-Ortiz, H., Rzhetskaya, M., Orozco, L., Raff, J.A., et al. (2019). Comparing signals of natural selection between three Indigenous North American populations. Proc. Natl. Acad. Sci. U. S. A. 116, 9312–9317.

31. Hider, J.L., Gittelman, R.M., Shah, T., Edwards, M., Rosenbloom, A., Akey, J.M., and Parra, E.J. (2013). Exploring signatures of positive selection in pigmentation candidate genes in populations of East Asian ancestry. BMC Evol. Biol. 13, 150.

32. Quillen, E.E., Bauchet, M., Bigham, A.W., Delgado-Burbano, M.E., Faust, F.X., Klimentidis, Y.C., Mao, X., Stoneking, M., and Shriver, M.D. (2012). OPRM1 and EGFR contribute to skin pigmentation differences between Indigenous Americans and Europeans. Hum. Genet. 131, 1073–1080.

33. Eaaswarkhanth, M., Xu, D., Flanagan, C., Rzhetskaya, M., Hayes, M.G., Blekhman, R., Jablonski, N.G., and Gokcumen, O. (2016). Atopic Dermatitis Susceptibility Variants in Filaggrin Hitchhike Hornerin Selective Sweep. Genome Biol. Evol. 8, 3240–3255.

34. Ojeda-Granados, C., Abondio, P., Setti, A., Sarno, S., Gnecchi-Ruscone, G.A., González-Orozco, E., De Fanti, S., Jiménez-Kaufmann, A., Rangel-Villalobos, H., Moreno-Estrada, A., et al. (2022). Dietary, Cultural, and Pathogens-Related Selective Pressures Shaped Differential Adaptive Evolution among Native Mexican Populations. Mol. Biol. Evol. 39.

35. Granka, J.M., Henn, B.M., Gignoux, C.R., Kidd, J.M., Bustamante, C.D., and Feldman, M.W. (2012). Limited evidence for classic selective sweeps in African populations. Genetics 192, 1049–1064.

36. Mathieson, I., and Terhorst, J. (2022). Direct detection of natural selection in Bronze Age Britain. bioRxiv, 2022.03.14.484330.

37. Olalde, I., Allentoft, M.E., Sánchez-Quinto, F., Santpere, G., Chiang, C.W.K., DeGiorgio, M., Prado-Martinez, J., Rodríguez, J.A., Rasmussen, S., Quilez, J., et al. (2014). Derived immune and ancestral pigmentation alleles in a 7,000-year-old Mesolithic European. Nature 507, 225–228.

38. Skoglund, P., Malmström, H., Omrak, A., Raghavan, M., Valdiosera, C., Günther, T., Hall, P., Tambets, K., Parik, J., Sjögren, K.-G., et al. (2014). Genomic Diversity and Admixture Differs for Stone-Age Scandinavian Foragers and Farmers. Science 344, 747–750.

39. Field, Y., Boyle, E.A., Telis, N., Gao, Z., Gaulton, K.J., Golan, D., Yengo, L., Rocheleau, G., Froguel, P., McCarthy, M.I., et al. (2016). Detection of human adaptation during the past 2000 years. Science 354, 760–764.

40. Abi-Rached, L., Jobin, M.J., Kulkarni, S., McWhinnie, A., Dalva, K., Gragert, L., Babrzadeh, F., Gharizadeh, B., Luo, M., Plummer, F.A., et al. (2011). The shaping of modern human immune systems by multiregional admixture with archaic humans. Science 334, 89–94.

41. Lenz, T.L. (2018). Adaptive value of novel MHC immune gene variants. Proc. Natl. Acad. Sci. U. S. A. 115, 1414–1416.

42. Phillips, K.P., Cable, J., Mohammed, R.S., Herdegen-Radwan, M., Raubic, J., Przesmycka, K.J., van Oosterhout, C., and Radwan, J. (2018). Immunogenetic novelty confers a selective advantage in host-pathogen coevolution. Proc. Natl. Acad. Sci. U. S. A. 115, 1552–1557.

43. Lazaridis, I., Nadel, D., Rollefson, G., Merrett, D.C., Rohland, N., Mallick, S., Fernandes, D., Novak, M., Gamarra, B., Sirak, K., et al. (2016). Genomic insights into the origin of farming in the ancient Near East. Nature 536, 419–424.

44. Skoglund, P., Malmström, H., Raghavan, M., Storå, J., Hall, P., Willerslev, E., Gilbert, M.T.P., Götherström, A., and Jakobsson, M. (2012). Origins and genetic legacy of Neolithic farmers and hunter-gatherers in Europe. Science 336, 466–469.

45. Olalde, I., Brace, S., Allentoft, M.E., Armit, I., Kristiansen, K., Booth, T., Rohland, N., Mallick, S., Szécsényi-Nagy, A., Mittnik, A., et al. (2018). The Beaker phenomenon and the genomic transformation of northwest Europe. Nature 555, 190–196.

46. Brace, S., Diekmann, Y., Booth, T.J., van Dorp, L., Faltyskova, Z., Rohland, N., Mallick, S., Olalde, I., Ferry, M., Michel, M., et al. (2019). Ancient genomes indicate population replacement in Early Neolithic Britain. Nat Ecol Evol 3, 765–771.

47. Günther, T., Valdiosera, C., Malmström, H., Ureña, I., Rodriguez-Varela, R., Sverrisdóttir, Ó.O., Daskalaki, E.A., Skoglund, P., Naidoo, T., Svensson, E.M., et al. (2015). Ancient genomes link early farmers from Atapuerca in Spain to modern-day Basques. Proceedings of the National Academy of Sciences 112, 11917–11922.

48. Günther, T., Malmström, H., Svensson, E.M., Omrak, A., Sánchez-Quinto, F., Kılınç, G.M., Krzewińska, M., Eriksson, G., Fraser, M., Edlund, H., et al. (2018). Population genomics of Mesolithic Scandinavia: Investigating early postglacial migration routes and high-latitude adaptation. PLoS Biol. 16, e2003703.

49. Yaka, R., Mapelli, I., Kaptan, D., Doğu, A., Chyleński, M., Erdal, Ö.D., Koptekin, D., Vural, K.B., Bayliss, A., Mazzucato, C., et al. (2021). Variable kinship patterns in Neolithic Anatolia revealed by ancient genomes. Curr. Biol. 31, 2455–2468.e18.

50. Coutinho, A., Günther, T., Munters, A.R., Svensson, E.M., Götherström, A., Storå, J., Malmström, H., and Jakobsson, M. (2020). The Neolithic Pitted Ware culture foragers were culturally but not genetically influenced by the Battle Axe culture herders. Am. J. Phys. Anthropol. 172, 638–649.

51. Sánchez-Quinto, F., Malmström, H., Fraser, M., Girdland-Flink, L., Svensson, E.M., Simões, L.G., George, R., Hollfelder, N., Burenhult, G., Noble, G., et al. (2019). Megalithic tombs in western and northern Neolithic Europe were linked to a kindred society. Proc. Natl. Acad. Sci. U. S. A. 116, 9469–9474.

52. Immel, A., Pierini, F., Rinne, C., Meadows, J., Barquera, R., Szolek, A., Susat, J., Böhme, L., Dose, J., Bonczarowska, J., et al. (2021). Genome-wide study of a Neolithic Wartberg grave community reveals distinct HLA variation and hunter-gatherer ancestry. Commun Biol 4, 113.

53. Rivollat, M., Jeong, C., Schiffels, S., Küçükkalıpçı, İ., Pemonge, M.-H., Rohrlach, A.B., Alt, K.W., Binder, D., Friederich, S., Ghesquière, E., et al. (2020). Ancient genome-wide DNA from France highlights the complexity of interactions between Mesolithic hunter-gatherers and Neolithic farmers. Sci Adv 6, eaaz5344.

54. Skourtanioti, E., Erdal, Y.S., Frangipane, M., Balossi Restelli, F., Yener, K.A., Pinnock, F., Matthiae, P., Özbal, R., Schoop, U.-D., Guliyev, F., et al. (2020). Genomic History of Neolithic to Bronze Age Anatolia, Northern Levant, and Southern Caucasus. Cell 181, 1158–1175.e28.

55. Furtwängler, A., Rohrlach, A.B., Lamnidis, T.C., Papac, L., Neumann, G.U., Siebke, I., Reiter, E., Steuri, N., Hald, J., Denaire, A., et al. (2020). Ancient genomes reveal social and genetic structure of Late Neolithic Switzerland. Nat. Commun. 11, 1915.

56. Villalba-Mouco, V., van de Loosdrecht, M.S., Posth, C., Mora, R., Martínez-Moreno, J., Rojo-Guerra, M., Salazar-García, D.C., Royo-Guillén, J.I., Kunst, M., Rougier, H., et al. (2019). Survival of Late Pleistocene Hunter-Gatherer Ancestry in the Iberian Peninsula. Curr. Biol. 29, 1169–1177.e7.

57. Mathieson, I., Alpaslan Roodenberg, S., Posth, C., Szécsényi-Nagy, A., Rohland, N., Mallick, S., Olalde, I., Broomandkhoshbacht, N., Cheronet, O., Fernandes, D., et al. (2017). The Genomic History Of Southeastern Europe. bioRxiv.

58. Mathieson, I., Alpaslan-Roodenberg, S., Posth, C., Szécsényi-Nagy, A., Rohland, N., Mallick, S., Olalde, I., Broomandkhoshbacht, N., Candilio, F., Cheronet, O., et al. (2018). The genomic history of southeastern Europe. Nature 555, 197–203.

59. Sánchez-Quinto, F., Schroeder, H., Ramirez, O., Ávila-Arcos, M.C., Pybus, M., Olalde, I., Velazquez, A.M.V., Marcos, M.E.P., Encinas, J.M.V., Bertranpetit, J., et al. (2012). Genomic Affinities of Two 7,000-Year-Old Iberian Hunter-Gatherers. Curr. Biol. 22, 1494–1499.

60. Olalde, I., Schroeder, H., Sandoval-Velasco, M., Vinner, L., Lobón, I., Ramirez, O., Civit, S., García Borja, P., Salazar-García, D.C., Talamo, S., et al. (2015). A Common Genetic Origin for Early Farmers from Mediterranean Cardial and Central European LBK Cultures. Mol. Biol. Evol. 32, 3132–3142.

61. Gamba, C., Jones, E.R., Teasdale, M.D., McLaughlin, R.L., Gonzalez-Fortes, G., Mattiangeli, V., Domboróczki, L., Kővári, I., Pap, I., Anders, A., et al. (2014). Genome flux and stasis in a five millennium transect of European prehistory. Nat. Commun. 5.

62. Saag, L., Varul, L., Scheib, C.L., Stenderup, J., Allentoft, M.E., Saag, L., Pagani, L., Reidla, M., Tambets, K., Metspalu, E., et al. (2017). Extensive Farming in Estonia Started through a Sex-Biased Migration from the Steppe. Curr. Biol. 27, 2185–2193 e6.

63. Allentoft, M.E., Sikora, M., Sjogren, K.-G., Rasmussen, S., Rasmussen, M., Stenderup, J., Damgaard, P.B., Schroeder, H., Ahlstrom, T., Vinner, L., et al. (2015). Population genomics of Bronze Age Eurasia. Nature 522, 167–172.

64. Broushaki, F., Thomas, M.G., Link, V., López, S., van Dorp, L., Kirsanow, K., Hofmanová, Z., Diekmann, Y., Cassidy, L.M., Díez-del-Molino, D., et al. (2016). Early Neolithic genomes from the eastern Fertile Crescent. Science 353, 499.

65. Jones, E.R., Zarina, G., Moiseyev, V., Lightfoot, E., Nigst, P.R., Manica, A., Pinhasi, R., and Bradley, D.G. (2017). The Neolithic Transition in the Baltic Was Not Driven by Admixture with Early European Farmers. Curr. Biol. 27, 576–582.

66. Marchi, N., Winkelbach, L., Schulz, I., Brami, M., Hofmanová, Z., Blöcher, J., Reyna-Blanco, C.S., Diekmann, Y., Thiéry, A., Kapopoulou, A., et al. (2022). The genomic origins of the world’s first farmers. Cell 185, 1842–1859.e18.

67. Delignette-Muller, M.L., Dutang, C., and Others (2015). fitdistrplus: An R package for fitting distributions. J. Stat. Softw. 64, 1–34.

